# KKL-35 exhibits potent antibiotic activity against *Legionella* species independently of trans-translation inhibition

**DOI:** 10.1101/135798

**Authors:** Romain Brunel, Ghislaine Descours, Isabelle Durieux, Patricia Doublet, Sophie Jarraud, Xavier Charpentier

## Abstract

Trans-translation is a ribosome rescue system that is ubiquitous in bacteria. Small molecules defining a new family of oxadiazole compounds that inhibit trans-translation have been found to have broad-spectrum antibiotic activity. We sought to determine the activity of KKL-35, a potent member of the oxadiazole family, against the human pathogen *Legionella pneumophila* and other related species that can also cause Legionnaires disease (LD). Consistent with the essential nature of trans-translation in *L. pneumophila*, KKL-35 inhibits growth of all tested strains at sub-micromolar concentrations. KKL-35 is also active against other LD-causing *Legionella* species. KKL-35 remains equally active against *L. pneumophila* mutants that have evolved resistance to macrolides. KKL-35 inhibits multiplication of *L. pneumophila* in human macrophages at several stages of infection. No resistant mutants could be obtained, even during extended and chronic exposure. Surprisingly, KKL-35 is not synergistic with other ribosome-targeting antibiotics and does not induce the filamentation phenotype observed in cells defective for trans-translation. Importantly, KKL-35 remains active against *L. pneumophila* mutants expressing an alternate ribosome-rescue system and lacking tmRNA, the essential component of trans-translation. These results indicate that the antibiotic activity of KKL-35 is not related to the specific inhibition of trans-translation and its mode of action remains to be identified. In conclusion, KKL-35 is an effective antibacterial agent against the intracellular pathogen *L. pneumophila* and with no detectable resistance. However, further studies are needed to better understand its mechanism of action and to assess further the potential of oxadiazoles in treatment.

## Introduction

*Legionella pneumophila* is a ubiquitous freshwater bacterium that infects a wide spectrum of environmental protozoans. Human-made systems such as sanitary water networks and air-cooling towers can disseminate contaminated water through aerosolization. Breathing microscopic droplets contaminated with *L. pneumophila* can lead to infection of alveolar macrophages and development of a life-threatening pneumonia called Legionnaire’s disease (LD) or Legionellosis. LD remains an important cause of both morbidity and mortality in Europe with over 6900 cases reported in 2014 (1). Guidelines for the management of LD recommend the use of macrolides (with a preference for azithromycin) or fluoroquinolones (levofloxacin/moxifloxacin) to treat the infection (2, 3). Despite a rapid diagnosis and the correct administration of antibiotics, death rate of LD is over 10% (4). *L. pneumophila* isolates are considered susceptible to macrolides and fluoroquinolones (5) but mutants resistant to both antibiotic families can be easily obtained *in vitro*, suggesting that resistant strains may emerge during treatment (6–8). Indeed, resistance to fluoroquinolones acquired in the course of a fluoroquinolone therapy has been recently reported (9, 10). New compounds active against *L. pneumophila* resistant to fluoroquinolones and macrolides or that could potentiate these existing treatments may improve the outcome of the disease.

Trans-translation has recently been proposed as a novel target for the development of a new class of antibiotics (11). Trans-translation is the primary bacterial mechanism to resolve ribosome stalling in bacteria (12–14). Ribosome stalling can be induced by translation of an mRNA lacking a stop codon (non-stop mRNA) or when ribosomes pause before the stop codon is read (*ie* due to ribosome-targeting antibiotics, rare sense codon stretches, lack of necessary tRNAs…). Ribosome stalling is a life-threatening issue in metabolically active bacteria (15, 16). Trans-translation is operated by a highly conserved nucleoprotein complex (17) encoded by two genes: *ssrA* encoding a highly expressed and structured RNA called tmRNA (18, 19), and *smpB* encoding a small protein involved in specific recognition and loading of tmRNA in stalled ribosomes (20–22). Once the complex is loaded into the free A-site of the stalled ribosome, translation resumes using the coding section of the tmRNA as template. This messenger section of tmRNA encodes a degradation tag that is appended to the unfinished polypeptide, targeting it to different proteases (23–25). The coding section of tmRNA ends with a stop codon, allowing normal termination of translation and dissociation of the ribosomal subunits. In addition, the tmRNA-SmpB complex interacts with RNAse R to degrade the faulty mRNA (26, 27). Thus, in addition to resolving ribosome stalling, the trans-translation system prevents the rise of further problems by promoting the degradation of both the problematic mRNA and the aborted polypeptide (28).

Alternative ribosome rescue systems have been identified in *Escherichia coli* and named ArfA and ArfB (Alternative rescue factors A and B) (29–31). Both ArfA and ArfB can partially complement the loss of trans-translation by promoting dissociation of the stalled ribosome but lack mechanisms to trigger degradation of the aborted polypeptide and faulty mRNA (12). These appear less conserved than the tmRNA-SmpB system (15). Trans-translation is essential in species lacking alternative mechanisms (16). In agreement with these observations, alternative ribosome-rescue systems are absent in members of the *Legionellaceae* genus and we indeed found that trans-translation is essential for *L. pneumophila* growth and infection of its cellular host (32). In *L. pneumophila*, expressing the alternate rescue factor ArfA from *E. coli* can compensate for the loss of trans-translation activity indicating that the ribosome-dissociating activity of the trans-translation system is the sole function required for viability (32). Because it is essential for viability in multiple pathogens, the trans-translation system has been proposed as a valid, yet-unexplored target for a new class of antibiotics (11).

A high-throughput screen using an *in vivo* assay of trans-translation recently identified a family of small molecule able to inhibit trans-translation at micromolar concentrations (33). One of the most active compounds, KKL-35, was found to exhibit a bactericidal activity against several pathogenic bacterial species in which trans-translation was known to be essential (33). KKL-35 and two related compounds KKL-10 and KKL-40 display antibiotic activity against the intracellular pathogen *Francisella tularensis* during infection of its host (34). However, the specificity of action of the molecules has not been confirmed in this species. The present study assessed KKL-35 activity against the intracellular pathogen, *L. pneumophila*. We report that KKL-35 exhibits potent antibiotic activity against *L. pneumophila* at very low concentrations and is able to stop bacterial multiplication in a model of infection of human macrophages. Yet, multiple evidence indicates that KKL-35 does not target trans-translation and, as such, its true target(s) in *L. pneumophila* remains to be identified.

## Results

### KKL-35 inhibits *Legionella* growth *in vitro*

Minimal inhibitory concentration (MIC) of KKL-35 were determined *in vitro* on five *L. pneumophila* strains and three non-*pneumophila* species causing LD using the broth microdilution method (Table 1). KKL-35 strongly inhibited growth of all tested species and was particularly potent against the species *Legionella pneumophila*, with all tested strains exhibiting a MIC around 0.04 mg/L. A time-kill assay on *L. pneumophila* str. Paris showed the bactericidal activity of KKL-35 with a decrease in viability at 24h after addition of KKL-35 at concentrations equal or higher than MIC (Figure 1A). At 72h following addition of KKL-35 at the MIC, the viable count was reduced by four orders of magnitude. Exposure to a half-MIC led to transient bacteriostatic activity for 48h, but then followed by growth suggesting that KKL-35 degrades and loses activity under those conditions. We also tested KKL-35 against twelve *L. pneumophila* mutants that were evolved from the Paris strain to become highly resistant to erythromycin and azithromycin (4000-fold increase in MIC) (8). The MIC of KKL-35 on these mutants was identical to that of the parent strain (0.04 mg/L) and thus unaffected by ribosomal mutations involved in macrolide resistance (23S rRNA, L4 and L22 proteins mutations). Interestingly, KKL-35 was poorly active in the conventional CYE solid medium for *L. pneumophila* with a MIC>10 mg/mL. A paper disk containing 100 µg of KKL-35 produces an inhibition zone of 7-8 mm in diameter (Figure 1B). To test the possibility that the agar-charcoal gelling base of CYE plates was reducing the activity of KKL-35, we substituted it by the guar gum gelling agent. On guar-gum plates (GYE), *L. pneumophila* form colonies exactly as on CYE plates (Figure 1C) but a disk of 100 µg of KKL-35 produces an inhibition zone of 40 mm in diameter (Figure 1B). KKL-35 at 0.02 g/L could already inhibit growth of inoculum with fewer than 10^7^ cfu and at 0.04 g/L (MIC in broth) no growth could be observed even with the highest inoculum (~10^8^ cfu) (Figure 1C). Thus, KKL-35 inhibits *L. pneumophila* growth both in liquid and solid medium at low concentration.

**Table 1.**
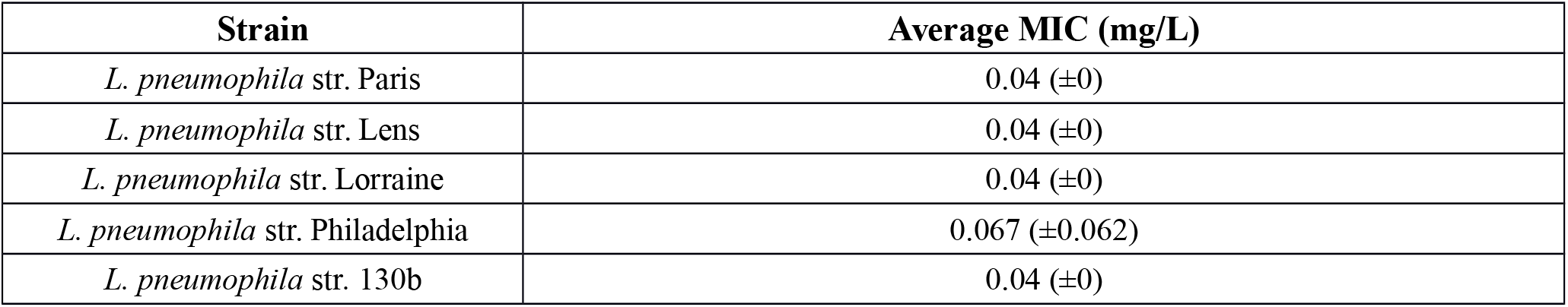
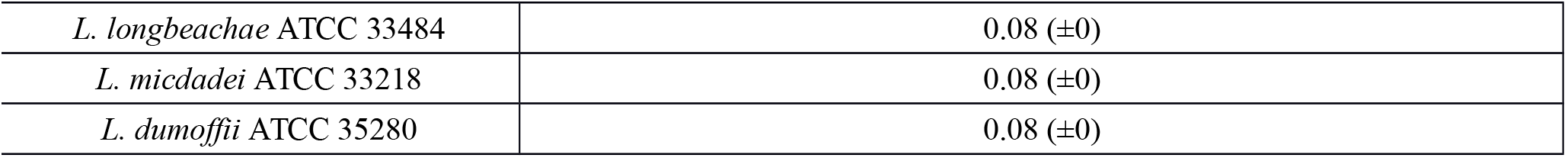
MIC of KKL-35 on several *Legionella* species *in vitro* (average and standard deviation from three independent determinations).

| Strain | Average MIC (mg/L) |
| --- | --- |
| *L. pneumophila* str. Paris | 0.04 (±0) |
| *L. pneumophila* str. Lens | 0.04 (±0) |
| *L. pneumophila* str. Lorraine | 0.04 (±0) |
| *L. pneumophila* str. Philadelphia | 0.067 (±0.062) |
| *L. pneumophila* str. 130b | 0.04 (±0) |
| *L. longbeachae* ATCC 33484 | 0.08 ($\pm 0$) |
| *L. micdadei* ATCC 33218 | 0.08 ($\pm 0$) |
| *L. dumoffii* ATCC 35280 | 0.08 ($\pm 0$) |

**Figure 1.**
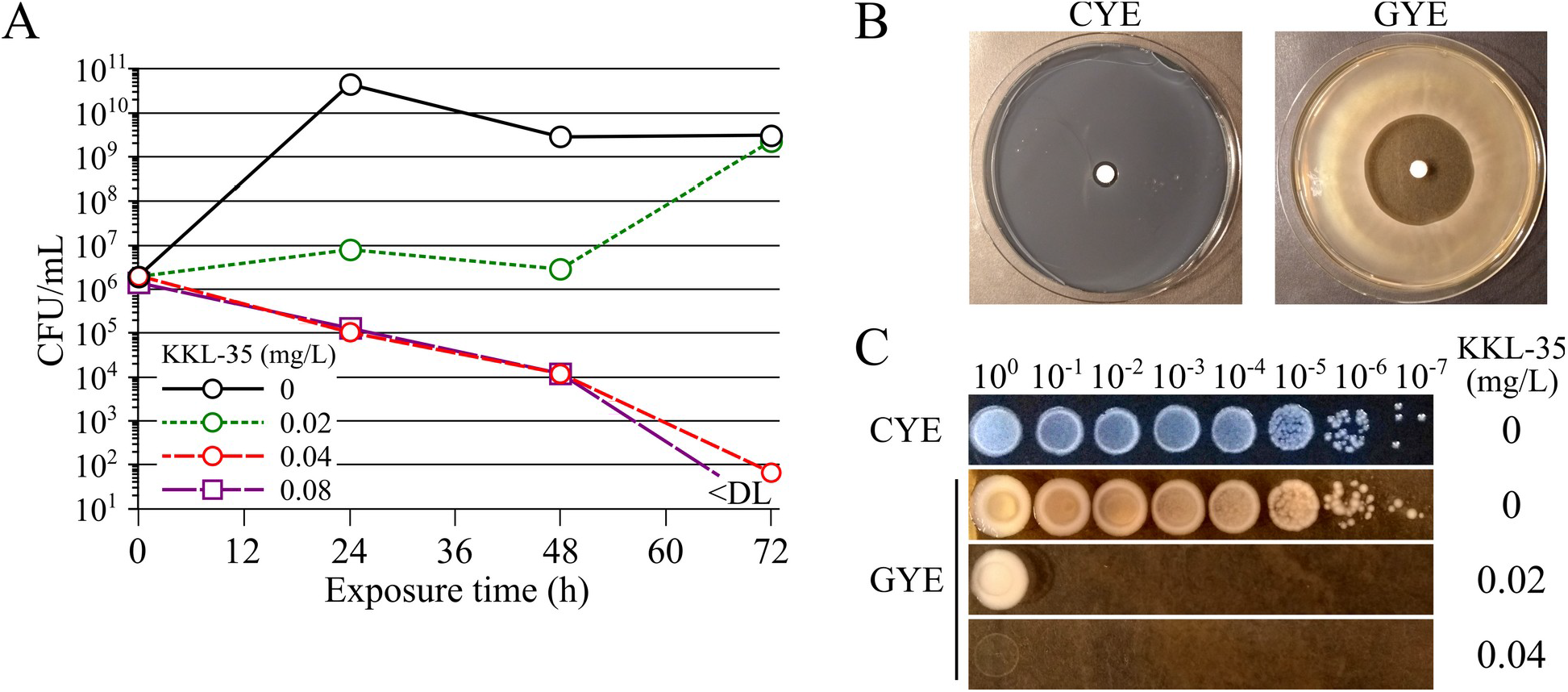
Antibiotic activity of KKL-35 on *L. pneumophila* in liquid and solid media. (A) Time-kill analysis of KKL-35 on *L. pneumophila* in liquid medium AYE. *L. pneumophila* strain Paris was resuspended in AYE medium at 3.10^6^ CFU/mL with a range of two-fold dilutions of KKL-35. Tubes were then incubated at 37°C. Every 24h, serial dilutions were plated on CYE agar and CFUs were counted. Presented data are average of triplicate samples. Black open circle, no KKL-35; green open circles, KKL-35 at 0.02 mg/L; red open circles KKL-35 at 0.04 mg/L; purple open squares KKL-35 at 0.08 mg/L. The presented data are representative of three experiments performed independently. (B) Antibiotic activity of KKL-35 in a solid medium disk diffusion assay. A paper disk containing 100 µg of KKL-35 was placed at the center of a CYE or GYE plate inoculated by flooding with a suspension of *L. pneumophila*. (C) Determination of the MIC of KKL-35 on GYE plates. Serial ten-fold dilutions of a culture of *L. pneumophila* in stationary phase (~5x10^9^ cfu/mL) were spotted (10 µl) on CYE (no KKL-35) and GYE plates containing increasing concentrations of KKL-35.

### KKL-35 inhibits intracellular growth of *L. pneumophila*

*L. pneumophila* can infect human macrophages and replicate extensively within a membrane-bound compartment until cell lysis. Two molecules, KKL-10 and KKL-40, structurally related to KKL-35 were found to be non-toxic to macrophages at concentrations up to 19 mg/L (34). Indeed, we found that KKL-35 was not toxic at 10 mg/L and even protected monocyte-derived macrophages from killing by *L. pneumophila* at a multiplicity of infection (MOI) of 10 (Figure 2A). In order to better characterize the inhibitory activity of KKL-35, we used a GFP-based time-resolved assay to follow the replication of GFP-expressing *L. pneumophila* in monocyte-derived macrophages (35). Within minutes of forced contact with macrophages, *L. pneumophila* is internalized in a vacuolar compartment that escapes fusion with lysosomes (36, 37). Addition of KKL-35 1h after infection, when bacteria are intracellular but not yet multiplying, prevented *L. pneumophila* replication at concentrations above 1 mg/L (Figure 2B). Moreover, when added at later timepoints (18 or 24h), when multiplication is ongoing, KKL-35 could inhibit replication at even lower concentrations (0.7 mg/L) (Figure 2B). This may indicate that either KKL-35 is more active against actively dividing cells or that the active fraction of KKL-35 gradually decreases over time. The ability of KKL-35 to completely halt replication was then compared to the activity of the macrolide erythromycin, a recommended treatment for LD. When added to actively replicating *L. pneumophila*, erythromycin begins to inhibit replication at 0.31 mg/L (Figure 2C). Each two-fold increase in concentrations further inhibit replication. A nearly complete and immediate inhibition of replication is obtained at a concentration 32 times higher than the first inhibitory concentration (10 mg/L). While KKL-35 begins to inhibit replication at a concentration of 0.62 mg/L, it can completely stop replication at a concentration only 8 times higher (5 mg/L) (Figure 2C). This indicate that KKL-35 may be more bactericidal than erythromycin. Altogether, the data show that KKL-35 inhibits replication of *L. pneumophila* within macrophages.

**Figure 2.**
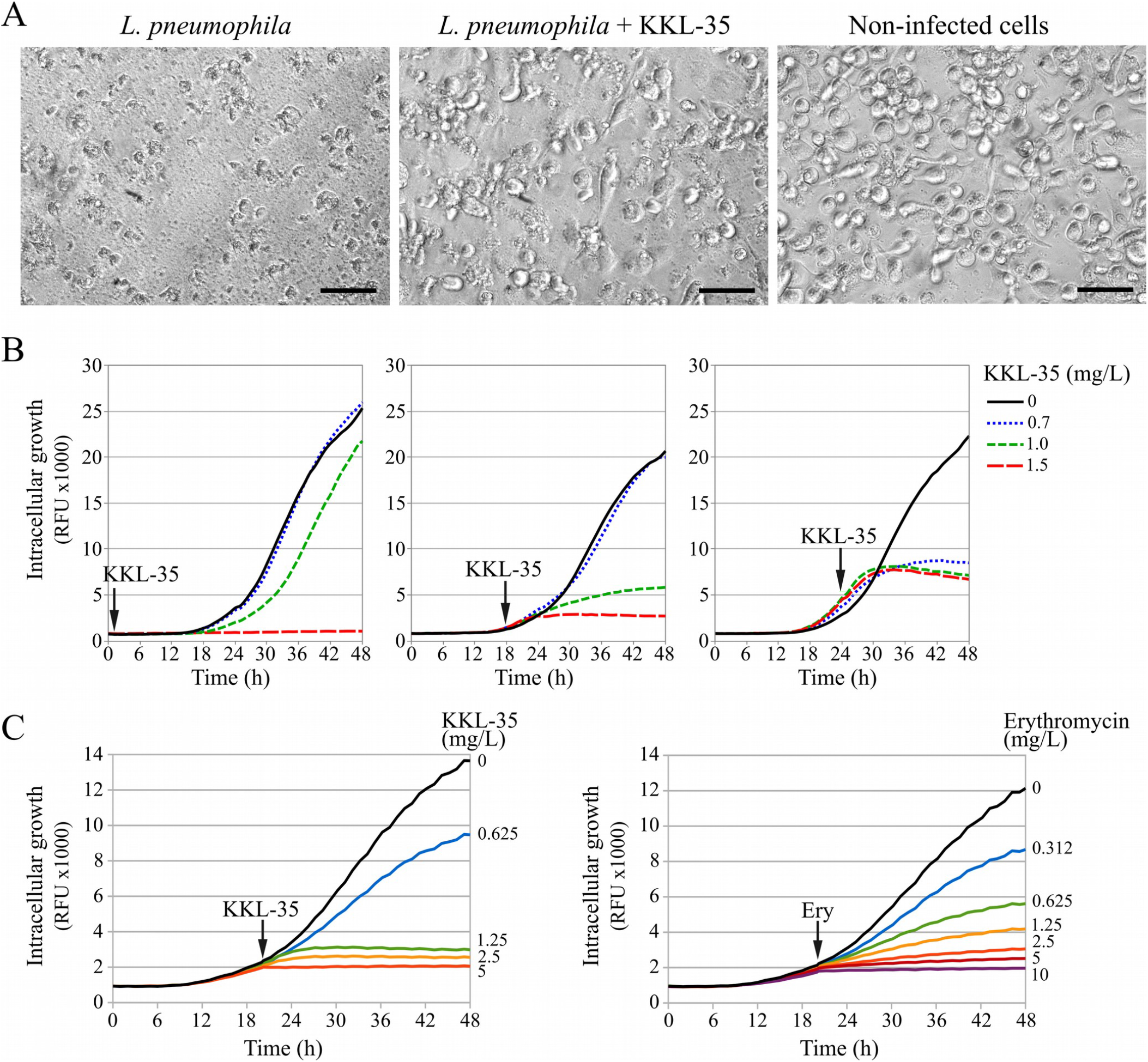
Activity of KKL-35 on *L. pneumophila* in an intracellular replication model. (A) Bright light microscopy imaging of U937-derived macrophages infected with *L. pneumophila* (MOI=10) for 72h in the presence of absence of KKL-35 at 10 mg/L. Black scale bar represents 50 µm. (B) Live monitoring of intracellular replication of GFP-producing *L. pneumophila* str. Paris carrying plasmid pX5 in U937-derived macrophages. KKL-35 was added at 1h, 18h or 24h post infection. GFP fluorescence levels were automatically monitored every hour for 48h. Black solid line, no KKL-35; blue short dashed line, KKL-35 at 0.7 mg/L; green intermediate dashed line, KKL-35 at 1 mg/L; red long dashed line, KKL-35 at 1.5 mg/L. RFU, relative fluorescence units. Data are average of three wells and representative of an experiment performed three times independently. (C) Comparison of the activity of KKL-35 and erythromycin on intracellular replication of *L. pneumophila*. KKL-35 and erythromycin were added at 20h post-infection. Data are average of three wells and representative of an experiment performed twice independently.

### KKL-35 does not induce phenotypes associated with loss of trans-translation

Lack of trans-translation increases the sensitivity to ribosome-targeting antibiotics in *E. coli* (38, 39) and in *L. pneumophila* (32). The *L. pneumophila* strains *ssrA*^ind^ carrying an IPTG-inducible allele of the tmRNA-encoding gene *ssrA* is unable to grow if IPTG is not supplied in the medium (32). Low levels of IPTG allow growth with artificially reduced levels of tmRNA, resulting in increased susceptibility to erythromycin and chloramphenicol (32). Complete lack of trans-translation may further increase the sensitivity of *L. pneumophila* to these antibiotics. Thus, we anticipated that KKL-35 could be synergistic with erythromycin and chloramphenicol. To determine a potential synergy we performed a chequerboard analysis (40). Interestingly, the MIC of erythromycin (0.125 mg/L) and chloramphenicol (1 mg/L) were not affected by KKL-35, indicating the absence of synergy (FICI=2). Thus, unlike the genetic alteration of trans-translation, KKL-35 does not potentiate activity of ribosome-targeting antibiotics. Another phenotype of *L. pneumophila* cells genetically deprived of tmRNA is extended filamentation, indicating that trans-translation is required for cell division (32). In contrast to *L. pneumophila* cells defective for trans-translation, *L. pneumophila* cells treated with KKL-35 at, below, or above the MIC, still display normal morphology (Figure 3A). The inability of KKL-35 to reproduce the phenotypes associated with loss of trans-translation suggests that its potent antibiotic activity is not primarily linked to inhibition of trans-translation.

**Figure 3.**
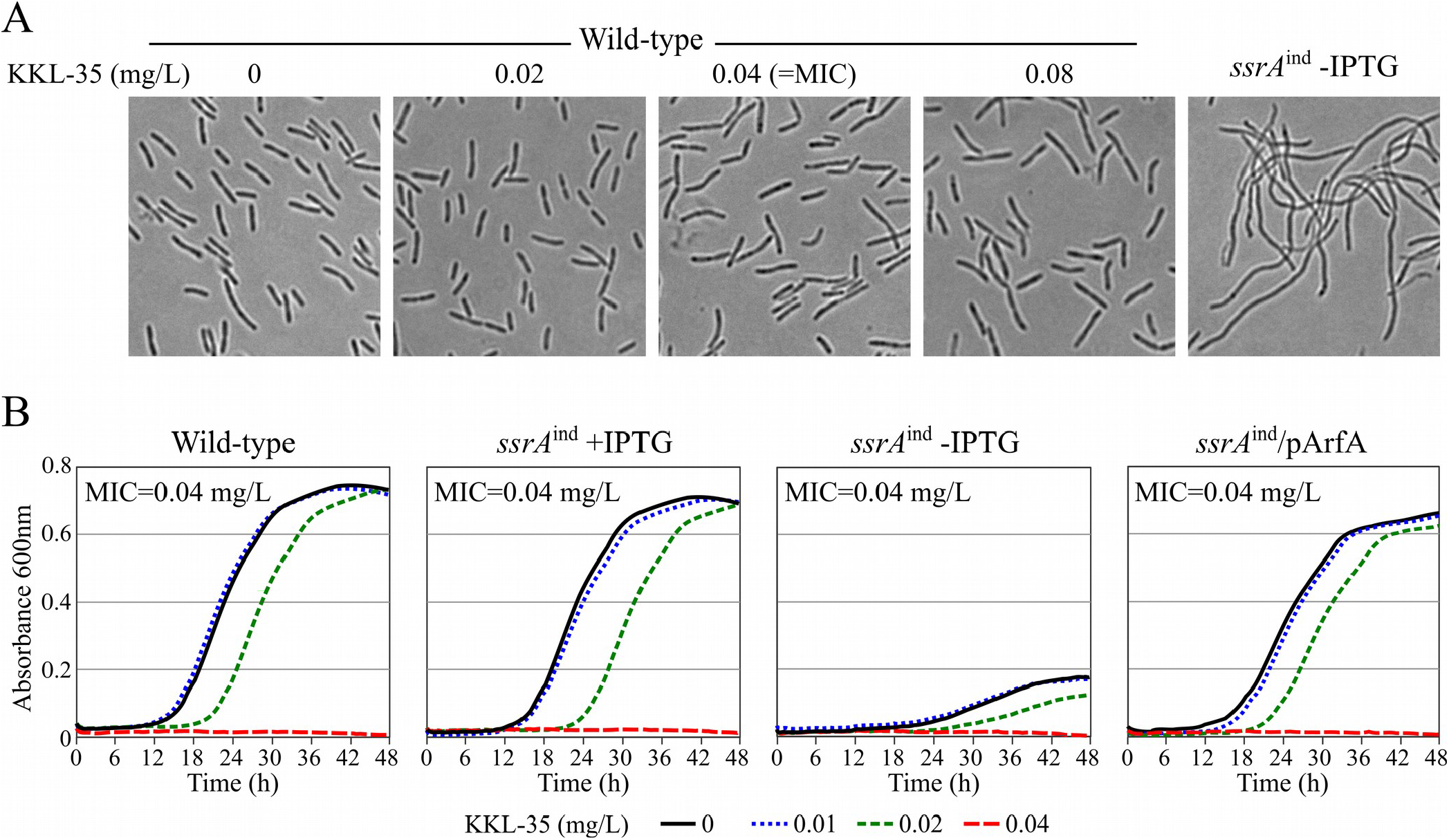
KKL-35 does not primarily target trans-translation in *L. pneumophila*. (A) Phase contrast light microscopy of wild-type *L. pneumophila* treated with KKL-35 for 24h and of the trans-translation deficient *ssrA*^ind^ mutant deprived of IPTG for 24h. (B) Activity of KKL-35 on *L. pneumophila* strains deficient for trans-translation. Representative growth curve of the wild-type, *ssrA*^ind^ mutant in the presence or absence of IPTG and of *ssrA*^ind^ mutant rescued by expression of the *E. coli* ArfA. MIC were determined three times independently on the basis of absorbance reading. Black solid line, no KKL-35; blue short dashed line, KKL-35 at 0.01 mg/L; green intermediate dashed line, KKL-35 at 0.02 mg/L; red long dashed line, KKL-35 at 0.04 mg/L.

### KKL-35 is equally active on *L. pneumophila* lacking trans-translation

To test whether the antibiotic activity of KKL-35 was linked to the inhibition of trans-translation, we tested KKL-35 on the *L. pneumophila* strains *ssrA*^ind^. When IPTG is supplied at high concentrations, tmRNA is expressed at near normal levels, and the strain grows like the wild-type strain. Expectedly, in the presence of IPTG this strain is equally sensitive to KKL-35 (MIC=0.04 mg/L) (Figure 3B). In the absence of IPTG, this strain is strongly impaired for growth. Yet, despite its low levels of tmRNA, the strain is not more sensitive to KKL-35. Ectopic expression of the alternate ribosome-rescue system ArfA from *E. coli* can restore growth of the *ssrA*^ind^ strain in the absence of IPTG. Under these conditions, the strain does not produce tmRNA and is therefore deficient for trans-translation (32). Despite not requiring trans-translation for growth, the MIC of KKL-35 on this strain remained identical to that on the wild-type strain (Figure 3B).

### *L. pneumophila* does not acquire resistance to KKL-35

*In vitro* selection of resistance is a common way to identify and characterize potential resistance determinants. Plating of large number of bacteria on solid medium containing antibiotic above the MIC often allows isolation of resistant mutants when resistance is conferred by a single mutation (i.e., rifampicin, streptomycin). Consistent with published data on *E. coli* (33), this strategy failed to produce mutants resistant to KKL-35 even when plating up to 10^10^ *L. pneumophila* cells on GYE plates containing KKL-35 at concentrations of 2 to 8 times the MIC (0.08-032 g/L). Occasionally, colonies could be obtained on GYE plates at the MIC (0.04 g/L) but the colonies could not grow again on freshly prepared GYE plates with same concentration of KKL-35 (data not shown). These colonies likely emerged because KKL-35 degrades over time. Continuous culture of a bacterial population in increasing concentrations of antibiotics represents an alternate approach when several mutations are required to confer resistance. In *L. pneumophila*, this method has been used to characterize the mutational path to resistance to fluoroquinolones and macrolides (7, 8). In agreement with previous reports, in two independent experiments, we here observed a 500-fold increase in the MIC of norfloxacin in only six passages (about 30 generations) (Figure 4). In contrast, no significant increase in the MIC of KKL-35 was obtained, even after 10 passages (over 60 generations) (Figure 4). Thus, in the tested experimental setup, *L. pneumophila* could not acquire resistance to KKL-35.

**Figure 4.**
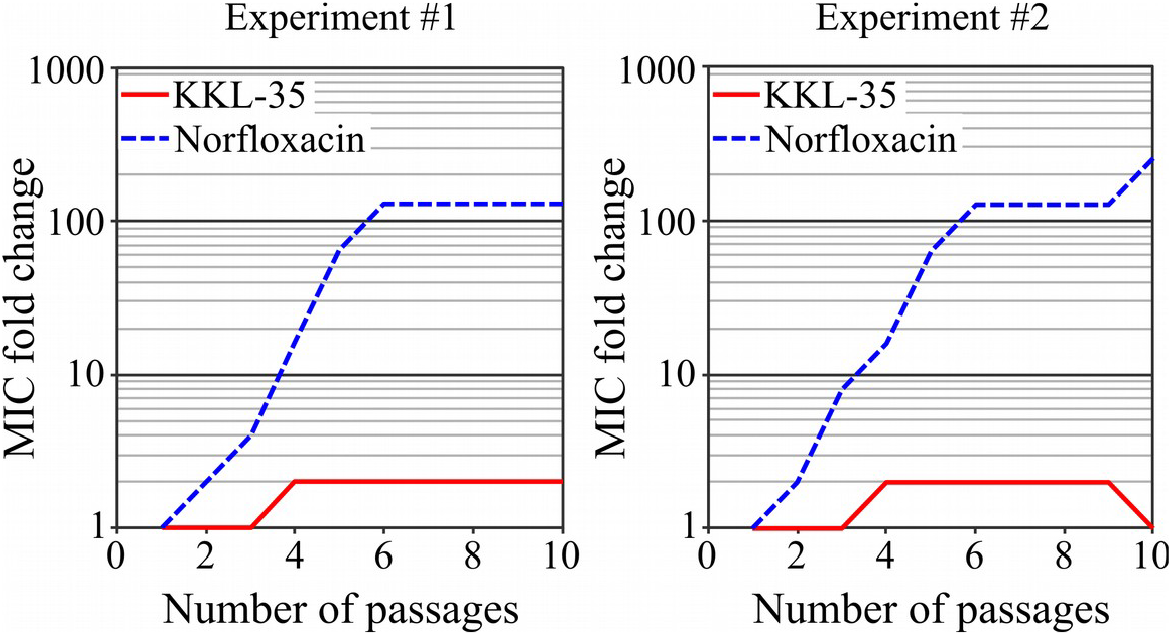
*L. pneumophila* does not acquire resistance to KKL-35. In two different experiments (several weeks apart), two different lineages were founded from *L. pneumophila* str. Paris and propagated by serial passages in the presence of KKL-35 (red solid line) or norfloxacin (blue dashed line). MIC was determined at each passage and presented relative to the initial MIC (norfloxacin: 0.25 mg/L, KKL-35: 0.04 mg/L).

## Materials and methods

### Strains, growth media and antibiotics used

*Strains used in this study included clinical isolates of L. pneumophila of strains Paris (CIP 107629), Lens (CIP 108286), Philadelphia-1, Lorraine (CIP 108729) and 130b, as well as isolates of L. longbeachae* ATCC 33484*, L. dumoffii* ATCC 35280*, L. micdadei* ATCC 33218. *L. pneumophila* str. Paris resistant to erythromycin or azithromycin were obtained from a previous work (8). *L. pneumophila* Paris was transformed with the plasmid pX5, a pMMB207C derivative harboring the *gfp+* gene under a strong constitutive promoter and used for live monitoring of intracellular multiplication by fluorescence reading. The tmRNA mutant strains *ssrA*^ind^ and *ssrA*^ind^/pArfA were previously described (32). ACES-Yeast Extract broth medium (AYE) was prepared with 10 g/L ACES (*N*-(2-acetamido)-2-aminoethanesulfonic acid), 12g/L yeast extract, 0.3 g/L iron III pyrophosphate and 0.5g/L L-cysteine. pH was adjusted at 6.9 with KOH and the solution was filter-sterilized and kept away from light, at 4°C. ACES-buffered charcoal yeast extract (CYE) plates were prepared by combining a two-fold concentrate of ACES and yeast extract (20 g/L each in the concentrate) with the same volume of and autoclaved solution of 30 g/L agar and 4 g/L charcoal (final concentrations 15 g/L agar and 2 g/L charcoal). The medium was then complemented with 0.25 g/L filtered iron III nitrate and 0.4 g/L L-cysteine and poured to produce CYE plates. Alternatively the agar-charcoal solution was substituted by an autoclaved guar gum solution in distilled water (2X concentrate 10 g/L, 5 g/L final) to produced Guar gum Yeast Extract plates (GYE). Unless indicated otherwise, cultures on CYE (or GYE) were incubated for 72h at 37°C in air, then patched onto CYE again for 24h to obtain fresh cultures before experiments were performed. When appropriate, chloramphenicol (5 µg/mL) was added to the medium. A stock solution of KKL-35 (Ambinter, Orléans, France) was prepared at 10 mM (3.2 g/L) in dimethylsulfoxide (DMSO) and stored at −20°C. KKL-35 was used at the highest at 10 mg/L (Figure 2A) on U937 cells or at a 1.5 mg/L on *L. pneumophila*, corresponding to a final concentration of DMSO of 0.3% and 0.05%, respectively.

### Time-kill assay, determination of MIC

For the time-kill assay, *L. pneumophila* strain Paris was resuspended in AYE medium at 3.10^6^ CFU/mL with a range of two-fold dilutions of KKL-35. Tubes were then incubated at 37°C in air, with shaking. Every 24h, serial dilutions were plated on CYE agar and CFUs were counted. For MIC determination, no CSLI guidelines are available for testing antibiotic susceptibility of *Legionella* strains. EUCAST guidelines were recently published but are based on the gradient strip test. KKL-35 strip tests are not commercially available, and we found the charcoal of CYE medium to seriously impede KKL-35 activity. Therefore, we used the previously described AYE broth microdilution method for MIC determination (41). Briefly, strains were resuspended in AYE medium and placed into the wells of a 96-well polystyrene plate and a range of twofold dilutions of KKL-35 was added to the cultures. The inoculum (10^6^ CFU/mL) was verified by plating and counting of serial dilutions of the cultures at the beginning of the experiment. The 96-well plate was sealed with a Breathe-Easy^®^ membrane (Sigma-Aldrich) to prevent evaporation and was incubated for 48h at 37°C in air with no agitation. At 48h, MICs were determined visually as the lowest concentrations inhibiting bacterial growth.

### Evaluation of synergistic activity

The chequerboard broth microdilution method was used to evaluate a possible synergistic activity between KKL-35 and chloramphenicol and erythromycin on *L. pneumophila* strain Paris. Bacteria were inoculated in AYE medium in a 96-well polystyrene plate containing a twofold range of concentration of KKL-35 in columns, crossing a range of another antibiotic in rows. The plate was then incubated for 48h in a Tecan Infinite M200Pro Reader at 37°C, with both agitation and absorbance reading at 600nm every 10 minutes. Growth value was defined as the highest absorbance reading recorded during the growth kinetic. Compared to the classic qualitative evaluation of growth by visual observation, this method allowed to obtain a quantitative measure of growth. Growth inhibition was defined as a maximal absorbance value <10% the value of the positive control. Fractional Inhibitory Concentration Index (FICI) were interpreted in the following way: FICI<=0.5 = synergy; FICI>4.0 = antagonism; FICI>0.5-4 = no interaction (40).

### Activity of KKL-35 on intracellular growth

U937 cells grown in RPMI 1640 containing 10% fetal calf serum (FCS) were differentiated into human macrophages by addition of phorbol 12-myristate 13-acetate (PMA) at 100 ng/mL, then seeded into 96-well polystyrene plates for 3 days (10^6^ cells/well). 4 hours before infection the medium was replaced with fresh medium + 10% FCS. *L. pneumophila* str. Paris was plated from a glycerol stock at −80°C onto CYE and incubated at 37°C in air for 72h, then plated again onto CYE plates for 24h to obtain a fresh culture. 4 hours before infection, bacteria were resuspended in RPMI 1640 and incubated at 37°C. Infection of macrophages was performed by replacing their medium by RPMI 1640 + 2% FCS containing *L. pneumophila* at a multiplicity of infection of 10. Plates were centrifuged 10 min at 1000 g then incubated at 37°C with 5% CO_2_ for 72h. Micrographs were taken with an inverted microscope (Nikon Eclipse TS100). Live monitoring of infection of U937 macrophages was performed as described above, except that the GFP-producing *L. pneumophila* str. Paris pX5 was used, and that the infection was performed in CO_2_-independent medium after differentiation, and was monitored by a Tecan Infinite M200Pro plate reader. The plate was incubated in the reader at 37°C and GFP fluorescence levels were automatically monitored every hour for 72h at an excitation wavelength of 470nm, and emission wavelength of 520nm.

### Selection of resistant mutants by serial passages

Two different lineages were founded from *L. pneumophila* str. Paris and propagated by serial passages in the presence of KKL-35 or norfloxacin, as previously described (7, 8). Briefly, a suspension of *L. pneumophila* str. Paris in AYE was added to a concentration of 10^8^ CFU/mL in a 24-well polystyrene plate with twofold KKL-35 or norfloxacin concentrations ranging from 0.5 times to 8 times the MIC that was determined for the parental strain (norfloxacin: 0.25 mg/L, KKL-35: 0.04 mg/L). Plates were sealed with a Breathe-Easy^®^ membrane (Sigma-Aldrich) and incubated for four days at 37°C in air without agitation, after which the minimum inhibitory concentration was noted for each antibiotic. Bacteria from the well with the highest antibiotic concentration in which growth was observable were transferred using a 1:40 dilution to a new plate containing twofold KKL-35 or norfloxacin concentrations ranging from 0.5 to 8 times the MIC of the previous cycle. Serial passages were repeated 10 times, and the experiment was performed twice independently.

## Discussion

We found KKL-35 to display a potent antibiotic activity against *L. pneumophila* with a MIC of 0.04 mg/L (0.125 µM). KKL-35 showed a significant bactericial activity and was found to retain a normal activity on different tested strains of erythromycin-resistant *L. pneumophila*. In addition, and in contrast to fluoroquinolones and macrolides, *L. pneumophila* did not develop resistance *in vitro*. Supporting a potential use in treatment of LD, KKL-35 could stop *L. pneumophila* from multiplying within monocyte-derived human macrophages. This indicates that KKL-35 is able to cross the biological membranes of the macrophage to reach intracellular *L. pneumophila*.

KKL-35, along with other oxadiazoles (KKL-10 and KKL-40), were identified in a high-throughput screen for inhibitors of trans-translation activity *in vitro* and were initially found to display antibiotic activity against several species. Oxadiazoles have since shown potent antibiotic activity against additional pathogens such as *Francisella tularensis*, *Bacillus anthracis* or *Mycobacterium tuberculosis* (34, 42, 43). The antibiotic activity of oxadiazoles on these pathogens was assumed to derive from inhibition of trans-translation but was not demonstrated. On *L. pneumophila*, several results led us to question the link between inhibition of trans-translation and antibiotic activity. First, no synergy was found between KKL-35 and ribosome-targeting antibiotics, whereas we previously found that a reduction in tmRNA levels led to an increased susceptibility of *L. pneumophila* to such antibiotics (32). Second, *L. pneumophila* cells treated with KKL-35 did not display the filamentation phenotype observed in cells lacking tmRNA. Most importantly, MICs were identical when KKL-35 was tested on a wild-type strain, on a growth-affected mutant expressing low levels tmRNA or on a mutant that does not require trans-translation because tmRNA was replaced by an alternative ribosome rescue system (ArfA from *E. coli*). Taken together our data indicate that trans-translation is not the primary target of KKL-35 in *L. pneumophila* and probably also in other bacteria. Indeed, similar results have been obtained in *E. coli* in which KKL-35 is equally effective on strains deficient for trans-translation (44). A novel double fluorescent reporter system for simultaneous and specific detection of trans-translation and proteolysis activities showed that KKL-35 has no direct effect on trans-translation (44). Altogether, the data challenge the initial report that KKL-35 is a trans-translation inhibitor, and that trans-translation is a promising antibiotic target (33). The molecular target of KKL-35 has not yet been identified but the impossibility to obtain resistant mutants indicate this target is not prone to support viable mutations. While this is a valuable property for an antibacterial agent, this hampered our efforts to identify the true target of KKL-35 in *L. pneumophila*. In conclusion, KKL-35 is an effective, broad-spectrum, antibacterial agent active against the intracellular pathogen *L. pneumophila*. However, further studies are needed to better understand its mechanism of action and to assess further the potential of oxadiazoles in treatment.

## Acknowledgements

We are grateful to Reynald Gillet for sharing unpublished results on the activity of KKL-35 and derivatives on *Escherichia coli*. We thank Floriane Nhoung and Noémie Fessy for technical assistance in MIC determination. RB is the recipient of a doctoral fellowship from the French Ministry of Higher Education and Research. This study has been funded by a Research Grant 2015 by the European Society of Clinical Microbiology and Infectious Diseases (ESCMID) awarded to XC.

